# Low ozone concentration and negative ions for rapid SARS-CoV-2 inactivation

**DOI:** 10.1101/2021.03.11.434968

**Authors:** Davide De Forni, Barbara Poddesu, Giulia Cugia, Giovanni Gallizia, Massimo La Licata, Julianna Lisziewicz, James Chafouleas, Franco Lori

## Abstract

Ozone is a powerful anti-bacterial, anti-fungal and anti-viral agent, yet exposure to high levels of ozone can pose risks to human/animal health and, in the long term, corrode certain objects. In order to overcome these risks, we evaluated the potential of using a relatively short exposure of a low concentration of ozone to disinfect an indoor environment in the absence of individuals and animals. ICON3 by O3ZONO/M2L, a new disinfection device generating both ozone and negative ions, was selected to assess the potential of this strategy to inactivate different viral isolates of SARS-CoV-2.

Tests under controlled laboratory conditions were performed in a system consisting of an ozone-proof airtight plastic box inside a biological safety cabinet, where suspensions of two strains of SARS-CoV-2 were exposed to ozone and negative ions and virucidal activity was measured by means of two complementary methodologies: viral replication capacity and viral titer determination.

These studies revealed that low concentration ozone (average 3.18 ppm after the peak) inactivated up to >99% of SARS-CoV-2 within 20 minutes of exposure. Under controlled conditions, similar ozone exposure was recreated with ICON3 in different volume rooms (15, 30, 60 m^3^) where a linear relationship was observed between the room volume and the time of continuous ozone/ions flow required to reach and maintain the desired ozone levels used in the laboratory studies.

These studies suggest that ICON3 may have the potential for use in the disinfection of SARS-CoV-2 in indoor environments in the absence of individuals and animals, under properly controlled and monitored safety conditions.

## Introduction

Coronavirus Disease 19 (COVID-19) was first identified in Wuhan (Hubei, China) in December of 2019, it has since been declared a pandemic by the World Health Organization (WHO) in March of 2020 [1, 2]. Following the initial cases in Wuhan, its pathogenic cause, Severe Acute Respiratory Syndrome Coronavirus 2 (SARS-CoV-2), has swept the world by March 2021 with over 120 million total cases and over 2.5 million deaths across 221 countries, areas, or territories. While recently approved vaccines might help to contain the viral spread, disinfection of the surrounding human environment are highly desirable. Social distancing, face masks, air ventilation, hand washing, ethanol spraying and chemical disinfection has been extensively used to decrease the reproduction rate (Rt) of this virus, a key measure of how fast the virus is growing, that is the average number of people who become infected by an infectious person. Unfortunately, these measures have not been sufficient to control virus transmission. There is a growing concern that part of the problem is constituted by the virus left in the public and private rooms, including in the air and over contaminated surfaces, similar to previous SARS infections [3, 4].

Ozone (O_3_) is a natural gas in the air that supports life on Earth by protecting all living forms against radiation. In the troposphere near the Earth's surface, the natural concentration of ozone is about 10 ppm, whereas it is absent at the surface or at low traces after some natural events like storms, lightning strikes and swells. Ozone is a powerful anti-bacterial, anti-fungal, and anti-viral agent [5–12]. It has been used for water purification and in healthcare facilities. Several ozone generators are available to emit ozone in high concentrations (e.g. up to several hundred ppm) to kill bacteria, fungi, and molds [5–7] and to inactivate viruses [8, 9]. Based on its mechanism of action, enveloped viruses, such as coronaviruses, are more sensitive to ozone than naked viruses [10–12].

The use of ozone generators to disinfect public and private rooms has been debated because of an uncertain risk-benefit balance and there is not a consensus among countries and governmental agencies regarding the ozone levels that might expose humans to risk and how ozone generators should be regulated. According to the US Environmental Protection Agency, exposure to ozone levels greater than 0.08 ppm for 8 hours or longer is unhealthy to humans. The major risks posed to human health by ozone are decrease in lung function, induction of inflammation and associated respiratory effects, including asthma and chronic obstructive pulmonary disease exacerbations [13]. In certain European countries ozone can be advertised exclusively as a sanitizer while it is presently being reviewed in Europe by the European Environmental Agency under the Biocidal Products Regulation of ECHA (European Chemicals Agency) for use as a biocide for surface disinfection [14]. Risks are not only represented by the potential harm to human/animal health but also by the potential damage to certain materials such as electric wire coating, rubber, and fabrics [15].

The benefit of ozone treatment against SARS-CoV-2 still needs to be carefully evaluated. For the disinfection of the indoor environment in the presence of individuals, exposure to continuous very-low concentration ozone was recently proposed. It was demonstrated that over 10 hours of treatment with 0.1 and 0.05 parts per million (ppm) ozone gas for 10 and 20 hours, respectively, that are the limits of permissible exposure for humans tolerated by the Japanese Society for Occupational Health and by and USA Food and Drug Administration, respectively, ozone is capable to decrease over 95% SARS-CoV-2 infectivity [16]. While the COVID-19 pandemic could potentially justify to use such a tolerated, continuous ozone exposure to protect individuals from infection, such an approach raises both safety and efficacy issues: on one hand concerns remain about the consequences of continuous ozone exposure on human health, without any reassurance that the ozone levels could be technically really kept below the threshold of toxicity; on the other hand, while it appears feasible to maintain such minimal levels of ozone in an experimental sealed room, it seems highly problematic to maintain a constant flow for a prolonged period of time in any environment where human beings are living and operating while air is continuously exchanged.

To disinfect the indoor environment a short exposure to a relatively low concentration of ozone in the absence of individuals/animals could instead be considered. ICON3, manufactured by O3ZONO (https://o3zono.it/home), is a new ozonizer that generates a flow of ozone pushed upwards along a vertical duct for an appropriate diffusion in the environment from a height of two meters. It is also an ionizer, generating a flow of negative ions. ICON3 can be programmed to emit ozone at a relatively low concentration (around an average of 3 ppm) together with ions (80 million/cm^3^). Exposure to such a low concentration of ozone for a relatively short time (approximately 20 minutes) would be expected to inactivate viruses, including SARS-CoV-2, without harming the environment and, in addition, it has been shown not to induce systemic oxidative stress in an animal model [17]. Although ions themselves are not expected to inactivate viruses, they remove ultrafine particles including aerosols of viruses from indoor air environments [18]. It is then anticipated that ion emission would add to the activity of the disinfection. Removing particles from the environment would also be beneficial, as a recent study demonstrated that exposure to fine particulate matter is associated with an increased incidence of COVID-19 [19].

Here it is demonstrated that a relatively low concentration ozone treatment (average 3 ppm after peak) with ICON3 inactivates up to >99% of SARS-CoV-2 within 20 minutes under controlled laboratory conditions inside an ozone-proof airtight plastic box under a biological safety cabinet. Moreover, similar ozone exposure parameters could be achieved under controlled conditions with the ICON3 in different sized rooms. When considering ICON3 disinfection of rooms, a strict control should be implemented, in the absence of individuals/animals and under reasonable surveillance, so that inactivation of SARS-CoV-2 can be attempted without adversely affecting the environment and human/animal health.

## Results

A new disinfection device, ICON3, was tested against SARS-CoV-2. ICON3 is an ozone and negative ions generator. It is built as a vertical structure, with the ozonizer/ionizer incorporated into the base of the device and with the ozone source placed at the top of a 2 meters high rod, allowing a constant ozone flow from top to bottom (Figure 1a). For testing in common rooms, the device was placed at one corner and the ozone was measured by means of a detector at different distances from the generator (see Materials and Methods).

**Figure 1.**
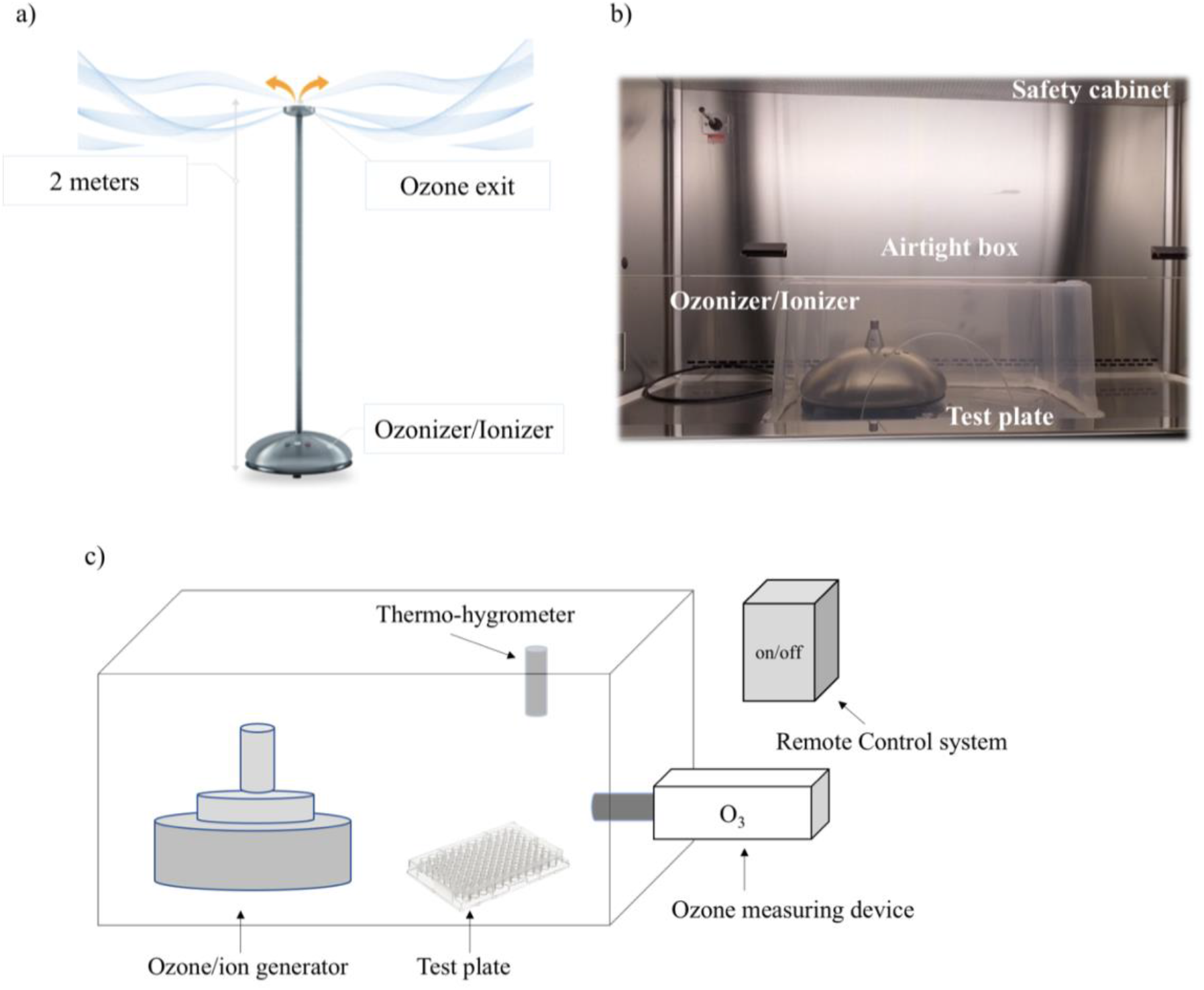
ICON3 ozone/negative ions generator. Figure Legend: ICON3 operating in a common room, with the ozone flow coming out of a 2 meters high rod from top to bottom (a); equipment for ozone gas administration in a plastic box under a biological safety cabinet (b); schematic representation of laboratory setup under biological safety cabinet (c).

For testing against SARS-CoV-2 in laboratory conditions, the rod was removed, and the device was placed inside an ozone-proof airtight plastic box (dimensions 57.3×39×25.7 cm; volume: 0.057 m^3^) inside a biological safety cabinet (Figure 1b, c). 0.5 μL drops of SARS-CoV-2 (2019-Nov/Italy-INMI1) suspension were deposited in a 96-well plate 15 cm away from the device. An ozone detector, i.e., a Pump Type Gas Detector, was positioned next to the plate. The ozone concentration in the plastic box was maintained among experiments between 5.44 and 1.47 ppm (mean 3.18 ± 1.5 ppm after peak) for 20 minutes after supplying a flow of ozone for approximately 1 second. Similar virus suspensions were exposed to air as control, then they were all collected to measure the replicative capacity of the virus (expressed as percentage of viral nucleoprotein, representing the replicating virus, compared to the control) in a Vero E6 cell line (see Methods). Exposure to ozone under these conditions inactivated SARS-CoV-2 by >99% (Table 1).

**Table 1.**
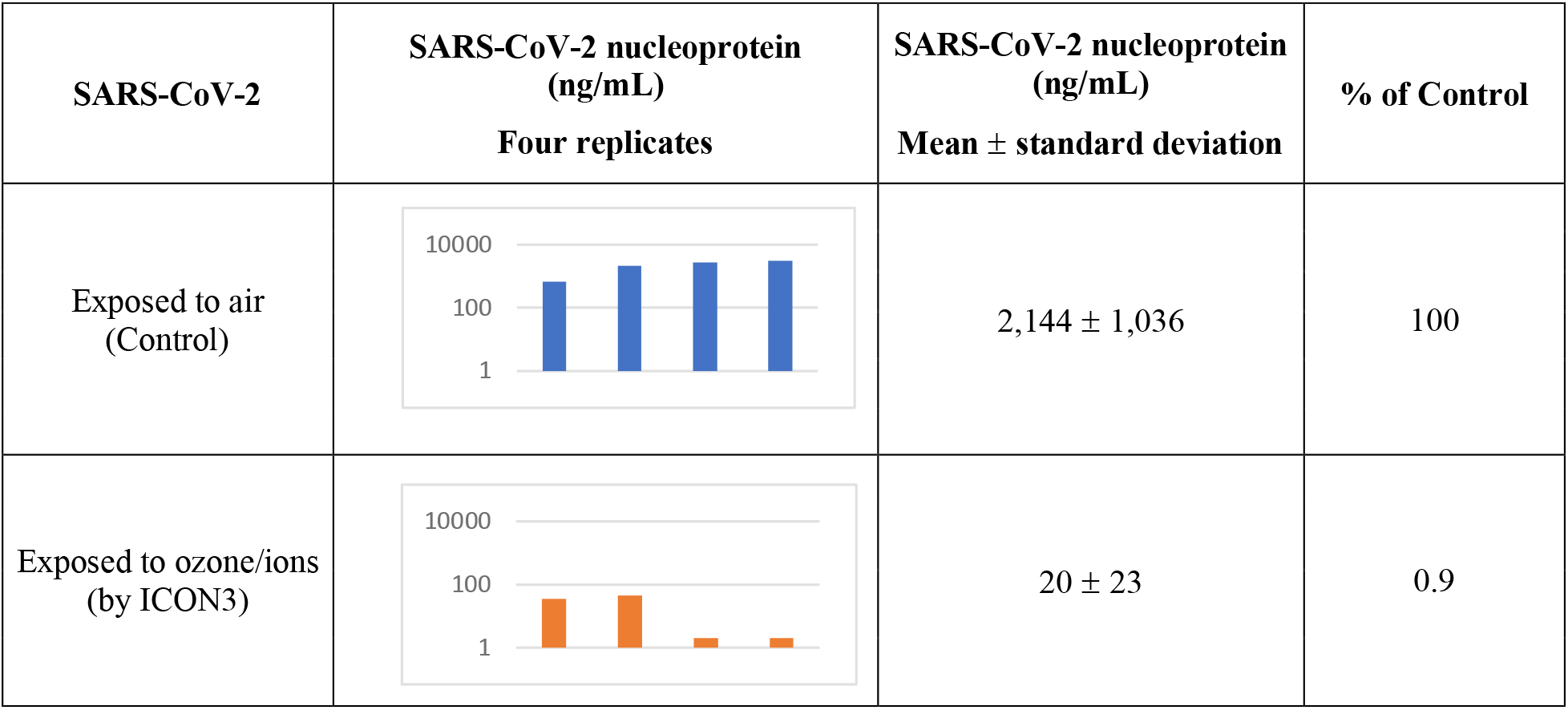
SARS-CoV-2 nucleoprotein reduction after exposure to ozone

| SARS-CoV-2 | SARS-CoV-2 nucleoprotein (ng/mL)<br>Four replicates | SARS-CoV-2 nucleoprotein (ng/mL)<br>Mean ± standard deviation | % of Control |
| --- | --- | --- | --- |
| Exposed to air<br>(Control) | | 2,144 ± 1,036 | 100 |
| Exposed to ozone/ions<br>(by ICON3) | | 20 ± 23 | 0.9 |

The experiment was repeated using a different readout, namely viral titer as opposed to a percentage of the active virus (see Materials and Methods). The results confirmed that SARS-CoV-2 (2019-nCov/Italy-INMI1 isolate) was inactivated by >99%. The experiment was performed with a different viral isolate (UNIBS-AP66: ERR4145453) that was also inactivated by >90% (Table 2). These results demonstrate that a low concentration of ozone can quickly and effectively inactivate different coronavirus isolates.

**Table 2.** Viral titer reduction of SARS-CoV-2 isolates after exposure to ozone

| SARS-CoV-2 | 2019-nCov/Italy-INMI1 |  | UNIBS-AP66: ERR414545 |  |
| --- | --- | --- | --- | --- |
|  | Viral titer (TCID<sub>50</sub>/mL) | Viral titer (% of control) | Viral titer (TCID<sub>50</sub>/mL) | Viral titer (% of control) |
| Exposed to air<br>(Control) | 7.94E+06 | 100 | 4.17E+06 | 100 |
| Exposed to ozone/ions<br>(by ICON3) | 3.98E+04 | 0.5 | 3.09E+05 | 7.4 |

Since droplets and aerosol of SARS-CoV-2 found in the air and surfaces that are responsible for transmission are quite small in size [20], it was investigated whether the size of the drops in which SARS-CoV-2 was resuspended would affect the results of the ozone treatment. During preliminary experiments it was determined that the smallest testable drop-size was 0.5 μL because smaller droplets quickly dried in the plate under the biological safety cabinet. 0.5 μL droplets are still considerably larger (by several logs) than the natural droplets originating during breathing, coughing, or sneezing [20]. The ICON3 device was used to expose different volumes of virus suspension (10, 3, 0.5 μL) to 5.44 - 1.47 ppm (mean 3.18 ppm) ozone for 20 minutes and measured the replicative capacity of the virus in Vero E6 cells (4 replicates each). Under such ozone exposure conditions SARS-CoV-2 was partially inactivated even when it was contained in very large droplets (Figure 2). Decreasing the droplet size further increased viral inactivation suggesting that SARS-CoV-2 contained in much smaller droplets originating from human coughing, sneezing or breathing might be completely inactivated. The results suggest that a relatively short exposure to the relatively low concentration of ozone generated by the ICON3 device under these controlled laboratory conditions might have the potential to disinfect the air and surfaces contaminated with SARS-CoV-2 naturally produced droplets.

**Figure 2.**
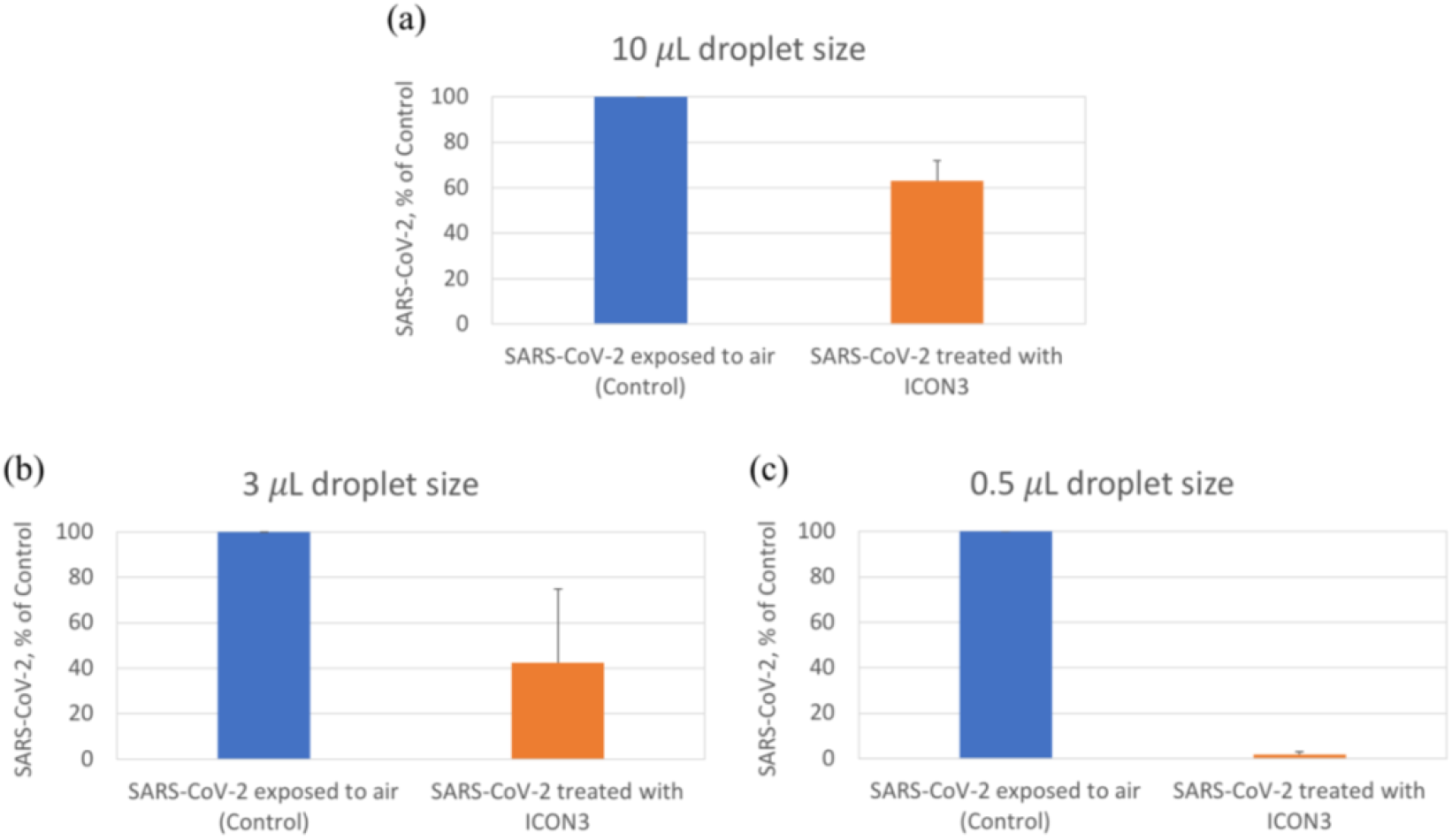
A low concentration of ozone inactivates SARS-CoV-2 in different size droplets. Figure Legend: ICON3 was used to expose SARS-CoV-2 to ozone/ions by keeping the ozone concentration between 5.44 - 1.47 ppm (mean 3.18 ppm) for 20 minutes (orange bars), compared to control (exposed to air, blue bars) and by varying the size of the droplets from 10 to 3 to 0.5 μL in Figures a, b and c, respectively. Replicative capacity is indicated as percentage viral nucleoprotein compared to control.

After demonstrating that a 20-minute exposure with a 3.18 ppm mean concentration of ozone can inactivate SARS-CoV-2 in the laboratory plastic box, it was investigated whether ICON3 could reproduce a similar ozone exposure in the most frequent sizes of an indoor environment, i.e. in three separate rooms of 15 m^3^, 30 m^3^, and 60 m^3^. In order to maintain similar ozone exposure conditions, the time for the device to remain ON (that is providing a continuous flow) had to be adapted according to the environment volume: only 1 second device ON time was needed in the laboratory cabinet (0.057 m^3^) while it was calculated that 4.5, 9, and 18 minutes would be required for the 15 m^3^ (2×2.5×3 m), 30 (3×3.3×3 m) m^3^, and 60 (4×5×3 m) m^3^ rooms, respectively. Under these precisely controlled conditions, exposure levels similar to that obtained in the laboratory plastic box were achieved in all three rooms (Figure 3 a, b, c, d). The Area Under the Curve (AUC), either Total, or Over Average or Under Average was in fact similar in all experimental conditions (Figure 3 e). The ozone decay time (time between switching off the ICON3 device and time to reach undetectable ozone levels) was also comparable, that is less than 45 minutes in each environment, more precisely 40, 37, 38 and 43 minutes in the laboratory cabinet, 15 m^3^, 30 m^3^, and 60 m^3^ rooms, respectively (Figure 3 e).

**Figure 3.**
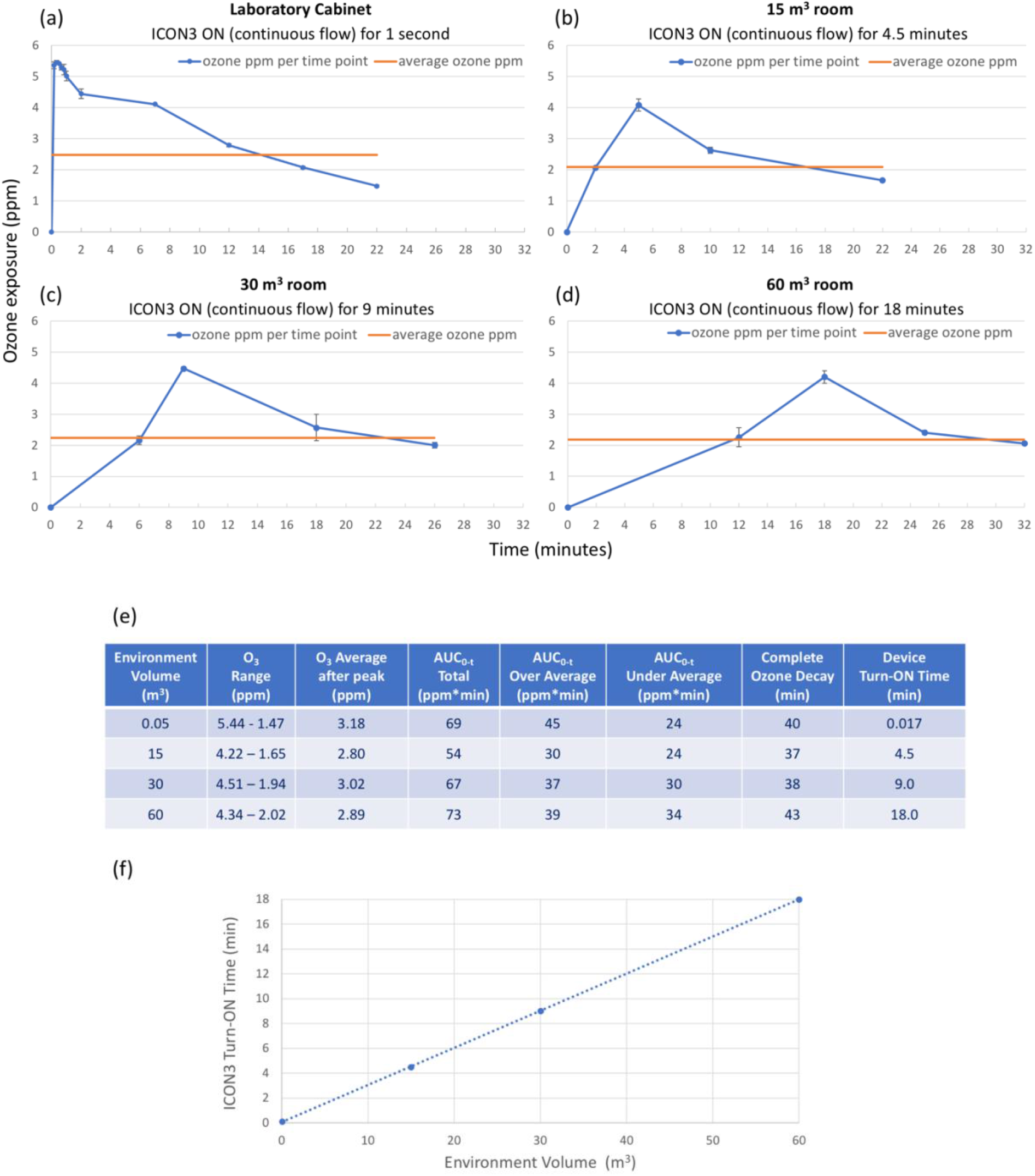
Keeping a similar ozone exposure in different volumes. Figure Legend: mean ozone exposure after a continuous flow maintained when ICON3 is turned on: for 1 second in a plastic box of 0.057 m^3^ under a biological safety cabinet (a); for 4.5 minutes in a 15 m^3^ room (b); for 9 minutes in a 30 m^3^ room (c); for 18 minutes in a 60 m^3^ room (d). Curves represent 2 repeated experiments in each ambient. A summary table indicating ozone exposure range, average, AUC values and ozone decay time (e); proportionality between environment volume and turn-ON time of ICON3, i.e., time of continuous ozone flow (f).

There was a linear relationship between the environment (room) volume and the time of continuous ICON3 flow required to obtain the desired ozone exposure (Figure 3 e, f). For example, doubling the room volume required a doubling in the time during which the ICON3 device remained switched ON in order to achieve the same overall ozone exposure to the one achieved in the laboratory cabinet.

## Discussion

Disinfection of indoor air environments, such as typical office or residential rooms, remains one of the main challenges in limiting transmission of viruses [21]. Currently these areas are disinfected by manual cleaning followed by chemical disinfection and/or ventilation. Disinfection is usually performed with alcohol-based or chlorinated solutions with or without ammonia. The safety of the person using these chemicals is essential. Cleaners must be trained to (i) wear adequate personal protective equipment such as gloves, medical masks, eye protection, and (ii) avoid combining disinfectants that could release gases causing respiratory irritations.

One of the main limitations of chemicals is that the efficiency of virus inactivation in the air and surfaces is somewhat problematic and it depends on human factors. For example, 1000 ppm hypochlorite could inactivate the vast majority of pathogens according to WHO [22], but several surfaces including furniture, mobile phones, computers, and other electronic devices, would be damaged by repeated spraying and infectious droplets would remain on these surfaces and mediate virus transmission. More importantly, cleaners use different chemicals, equipment, procedures, and their performances are not comparable nor reproducible. High concentrations (over 350 ppm) of ozone are used to inactivate SARS-CoV-2 in healthcare facilities, however, such high ozone concentrations are not commonly used in public offices and households because of safety concerns.

To overcome the limitations of chemical disinfection and high concentration of ozone the ICON3 device was calibrated to emit low concentration ozone and ions to effectively and reproducibly inactivate SARS-CoV-2. Although ions produced by the ICON3 are not expected to directly inactivate SARS-CoV-2 they might contribute to precipitate virus-containing droplets and decrease the levels of small particles that are associated with increased transmission of COVID-19 [19]. The objective of this study was to keep the ozone concentration around 3 ppm for a relatively short time (i.e., 20 minutes), conditions expected to not damage the surrounding material and be applicable for use in the absence of humans/animals [15]. Under such controlled conditions up to >99% SARS-CoV-2 was inactivated in a plastic box under a biological safety cabinet in the laboratory. It was demonstrated that similar ozone exposure parameters could be achieved in several different sized rooms where the time to reach the desired ozone concentration in a given room was linearly correlated to the room volume, i.e., doubling the volume of the room required a doubling of the time of continuous ozone/ions flow. Ozone decay time was similar and independent of the environment volume.

Disinfection by the ICON3 device offers distinctive features compared to other currently used methods: (i) experimental evidence has been provided here that ozone and ions emitted by the ICON3 device inactivate different SARS-CoV-2 strains up to >99% under controlled conditions; (ii) ozone is expected to penetrate everywhere and to inactivate viruses not only on selected surfaces that a cleaner disinfects but on all the surfaces present in the room as well as in the air; (iii) under the controlled conditions of this study a linear relationship between time of device ON and room volume provides a means to program the device to achieve the ozone exposure parameters obtained in the laboratory box study; (iv) since ozone quickly degrades to oxygen, after the appropriate ozone decay time, individuals returning to the ozone disinfected rooms would not be harmed; (v) furniture and equipment in the room should not be damaged; (vi) ozone would not contaminate the environment with chemical waste.

Several SARS-CoV-2 variants are now circulating globally, most notably: in the United Kingdom (UK), a new variant of SARS-CoV-2 (known as 20I/501Y.V1, VOC 202012/01, or B.1.1.7); in South Africa, another variant of SARS-CoV-2 (known as 20H/501Y.V2 or B.1.351); in Brazil, a variant of SARS-CoV-2 (known as P.1) [23]. All these variants present mutations in the receptor binding domain of the spike protein and there is some evidence to indicate that for example one of the spike protein mutations (E484K, shared by B.1.351 and P.1 variants) may affect neutralization by some polyclonal and monoclonal antibodies [24, 25].

Ozone virucidal effects are linked to its ability to break apart lipid molecules with multiple bonds, in fact, enveloped viruses are usually more sensitive to physical–chemical challenges than naked ones. Ozone can also interact with proteins, carbohydrates, and nucleic acids [26–28]. The ozone virucidal effect is therefore largely independent from the genome of SARS-CoV-2 and is expected to be effective against multiple existing and newly emerging variants. Consistent with this expectation, we and others [16] were able to achieve inactivation of three different SARS-CoV-2 variants to a similar extent (over 90%).

In this study, SARS-CoV-2 was inactivated to a large extent and in a relatively short time by using ICON3 under controlled laboratory conditions. This new ozone and ion emitting device, developed for use in public rooms and common living areas, may have the potential for use in the disinfection of SARS-CoV-2 in indoor environments in the absence of individuals and animals, under properly controlled and monitored safety conditions.

## Materials and Methods

### Viral isolates

Human 2019-nCoV strain 2019-nCoV/Italy-INMI1 was isolated in Italy (ex-China) from a sample collected on January 29, 2020, from the Istituto Lazzaro Spallanzani, Rome, Italy. A second strain, namely SARS-CoV-2-UNIBS-AP66: ERR4145453 [29, 30] was obtained from the University of Brescia, Italy.

### ICON3 device

ICON3 is an ozonizer/ionizer manufactured by O3ZONO [31]. The device generates a flow of ozone of 5000 mg/hour, with an exit volume of 1,200 liters/hour. It is also an ionizer, generating a flow of negative ions (80 million/cm^3^/3 minutes according to manufacturer’s specifications). The device consists of a base (diameter 30 cm, height 15 cm), containing the ozonizer and the ionizer device, and a detachable 2-meter-high rod, at the top from which ozone flows (Figure 1).

### Treatment of viral suspensions with ICON3

Indicated volumes of SARS-CoV-2 suspensions were exposed to ozone and negative ions produced by the ICON3 device. The treatment process was conducted inside a plastic box (dimensions 57.3×39×25.7 cm; volume: 0.057 m^3^), where the ICON3 machine was placed, together with the viral suspension in a 96-well plate. The box was maintained under a biosafety cabinet for safety reasons. Quantity of ozone was monitored during treatment with a Pump Type Gas Detector (detection range 0-50 ppm, resolution 0.01 ppm). The device was turned on for 1 second allowing ozone to reach a maximum peak of 5.44 ppm. Contact time was 20 minutes. The test temperature was 21°C C 1°C. Relative humidity in the cabinet was 50 C 1%. An identical volume of viral suspension exposed to the air for the same contact time was used as control.

### Determination of SARS-CoV-2 replicative capacity

The replicative capacity of treated viral suspensions was measured in a cell model. Vero E6 cells (kidney epithelial cells from African green monkey, ATCC CRL-1586) were maintained at their optimal density based on the ATCC datasheet. On Day 1 of the experiment, cells were transferred in a 96-well plate (10.000 cells per well). On Day 2 cells were infected with the different viral suspensions (treated and not treated with ozone and negative ions) in quadruplicate wells at a multiplicity of infection (m.o.i.) of 0.01. After 3 additional days, supernatants were collected, and an ELISA assay (SARS-CoV-2 Nucleocapsid Detection ELISA Kit, Sino Biological) was performed to measure produced virus through the quantification of the viral NP nucleoprotein (a measure of viral replication capacity).

### Determination of SARS-CoV-2 viral titer

An *in vitro* system was employed to determine the viral titer of both SARS-CoV-2 treated with ozone and negative ions as well as of SARS-CoV-2 exposed to air. Vero E6 cells (kidney epithelial cells from African green monkey, ATCC CRL-1586) were maintained at their optimal density based on the ATCC datasheet. On Day 1 of the experiment, cells were transferred in a 96-well plate (10.000 cells per well). On Day 2 cells were infected with the serial viral dilutions (10^-2^, 10^-^ 3, 10^-4^...) in 6-well replicates for each condition. After 3 additional days, supernatants were collected and an ELISA assay (SARS-CoV-2 Nucleocapsid Detection ELISA Kit, Sino Biological) was performed to determine infection or not of each test well, through the detection of the viral NP nucleoprotein. Data were then used to determine the titer of treated and control virus according to Reed and Muench method.

### Measuring ozone in the rooms and in the laboratory

Quantity of ozone was monitored with a Pump Type Gas Detector in three separate rooms of 15 m^3^, 30 m^3^, and 60 m^3^ and the detector was positioned at 1-, 2- and 3-meters distance from the ozone/ions generator, respectively, and at 120 cm height. The measurements were repeated two times within the same room and presented as Average and Standard Deviation at each time point. For testing against SARS-CoV-2 in laboratory conditions, the rod was removed, and the device was placed inside an ozone-proof airtight plastic box (dimensions 57.3×39×25.7 cm; volume: 0.057 m^3^) inside a biological safety cabinet. The ozone decay time was calculated in similar, separate experiments comparing the three different rooms and the plastic box under the biological safety cabinet.

## Acknowledgments

This publication was supported by the European Virus Archive GLOBAL (EVA-GLOBAL) project that has received funding from the European Union’s Horizon 2020 research and innovation program under grant agreement No 87102.

We thank Arnaldo Caruso and Francesca Caccuri, from the Microbiology and Virology Section, Molecular Medicine Department, University of Brescia, Italy for providing viral strain SARS-CoV-2-UNIBS-AP66: ERR4145453. We also thank Sjlva Petrocchi for editorial assistance.

We also thank Sjlva Petrocchi for her comments and revisions.

